# Prediction of quality-control degradation signals in yeast proteins

**DOI:** 10.1101/2022.04.06.487301

**Authors:** Kristoffer E. Johansson, Bayan Mashahreh, Rasmus Hartmann-Petersen, Tommer Ravid, Kresten Lindorff-Larsen

## Abstract

Effective proteome homeostasis is key to cellular and organismal survival, and cells therefore contain efficient quality control systems to monitor and remove potentially toxic misfolded proteins. Such general protein quality control to a large extent relies on the efficient and robust delivery of misfolded or unfolded proteins to the ubiquitin-proteasome system. This is achieved via recognition of so-called degradation motifs—degrons—that are assumed to become exposed as a result of protein misfolding. Despite their importance, the nature and sequence properties of quality-control degrons remain elusive. Here, we have used data from a yeast-based screen of 23,600 17-residue peptides to build a predictor of quality-control degrons. The resulting model, QCDPred (Quality Control Degron Prediction), achieves good accuracy using only the sequence composition of the peptides as input. Our analysis reveals that strong degrons are enriched in hydrophobic amino acids and depleted in negatively charged amino acids, in line with the expectation that they are buried in natively folded proteins. We applied QCDPred to the yeast proteome, enabling us to analyse more widely the potential effects of degrons. As an example, we show a correlation between cellular abundance and degron potential in disordered regions of proteins. Together with recent results on membrane proteins, our work suggest that the recognition of exposed hydrophobic residues is a key and generic mechanism for proteome homeostasis. QCDPred is freely available as open source code and via a web interface.

## Introduction

Most proteins are required to fold into their native three-dimensional structures to carry out their cellular functions. Mutations or environmental stress conditions may, however, lead to partial unfolding or misfolding of these natively folded proteins, or to disassembly of complexes. Within the cell, such structurally unstable or misfolded proteins may form aggregates and often display proteotoxicity via poorly understood mechanisms. Cells are therefore equipped with an effective protein quality control (PQC) system that catalyses the refolding or degradation of aberrant proteins to sustain proteostasis ^1–3^. The degradative route of the eukaryotic PQC system often relies on the ubiquitin-proteasome system to clear the cell of non-native proteins. Here, E3 ubiquitin-protein ligases, frequently assisted by molecular chaperones, first conjugate the non-native proteins to ubiquitin-chains, which in turn target the proteins for proteasomal degradation ^4–8^.

Many E3s are involved in specific cellular functions and are highly specific enzymes ^9, 10^. As virtually all cellular proteins may become misfolded, it is, however, essential that the PQC-linked degradation system is not too specific so that it can degrade a wide range of misfolded substrates, but simultaneously leaves native proteins intact. The exact substrate preferences of the PQC E3s are largely unknown, and accordingly, the discriminating feature(s) of misfolded PQC targets are unknown and presently not possible to predict. E3s recognize regions in target proteins termed degradation signals or degrons ^11, 12^. The so-called primary degron interacts—directly or indirectly via adaptor proteins—with the E3, and thus determines specificity. In addition, a nearby lysine residue that acts as the acceptor site for ubiquitylation, the so-called secondary degron, is also required. Finally, a partially unfolded or disordered region in the substrate protein, termed the tertiary degron, is required for the proteasome to commence threading the protein into the particle for degradation ^13^. We here focus on the properties of primary PQC degrons that determine the specificity in PQC-linked proteasomal degradation. Despite several efforts ^14–16^, the discerning features of the PQC degrons remain rather enigmatic. These degrons are, however, expected to include hydrophobic areas that are buried in the native protein but become exposed upon unfolding or misfolding.

Previously, Geffen et al. used a Ura3-GFP reporter system to map large numbers of degradation signals in *Saccharomyces cerevisiae* ^15^. These experiments revealed a large breadth in the sequences that cause degradation without any obvious individual motifs. More recently, we used the same reporter system to show that Hsp70-binding motifs may function as PQC degrons ^17^. Specifically, we found that inserting known Hsp70 binding motifs into the reporter system led to Hsp70-dependent degradation.

In a new set of experiments, we extended these ideas to map more systematically degrons in *Saccharomyces cerevisiae* ^18^. Specifically, we used a fluorescence-based reporter system coupled to fluorescence-activated cell sorting (FACS) and sequencing to study the degradation of 23,600 17-residue long peptides. Here, we use this data to study the properties of the degrons and build a prediction method for PQC degrons. Our analysis confirms our previous observation that groups of hydrophobic residues—often buried inside the core of folded proteins—can act as strong degrons when exposed to the PQC system. We also show that negatively charged residues counteract the degradation signal, potentially explaining why these are common in many disordered proteins. Finally, we show that the presence of PQC degrons in disordered regions of proteins may influence their cellular abundance.

## Results and Discussion

We aimed to develop a prediction algorithm to determine which protein sequences, when exposed to the cytosolic PQC system, can be recognized and targeted for degradation. As input to our model, we used data from an assay in which 23,600 17-residues long peptides from the yeast proteome had been fused to GFP in a reporter construct ^18^. In this construct, peptides with strong degron properties cause degradation of GFP which, when normalized by an expression control, can be distinguished from non-degron peptides using FACS and quantified by sequencing. Details and validation of the experiments are described elsewhere ^18^ and briefly summarized in the methods section.

Using the sequencing counts to quantify which peptides lead to low or high levels of GFP, we separate the assayed peptides into six bins based on their protein stability index (PSI) (Fig. 1A). Here a high PSI score indicates peptides associated with high GFP fluorescence (non-degrons), and a low PSI score with peptides that cause decreased GFP fluorescence (degrons). The value of the PSI score between one and four reflects the average position of a peptide among the four bins from the FACS analysis. As discussed below and elsewhere ^18^, the decreased GFP signal is in most cases due to proteasomal degradation resulting from the fusion with the peptide. We divided the peptides into six bins in units of 0.5 PSI score, and find that the amino acid composition of the peptides in each of the six bins shows a clear trend towards increased hydrophobicity in peptides that are associated with low GFP signals (likely degrons) and negatively charged side chains in peptides that do not act as degrons (Fig. 1A and Fig. S1). Most proteins have both degron and non-degron sequences and have, on average, tiles in 4.7 of the 6 bins.

**Figure 1.**
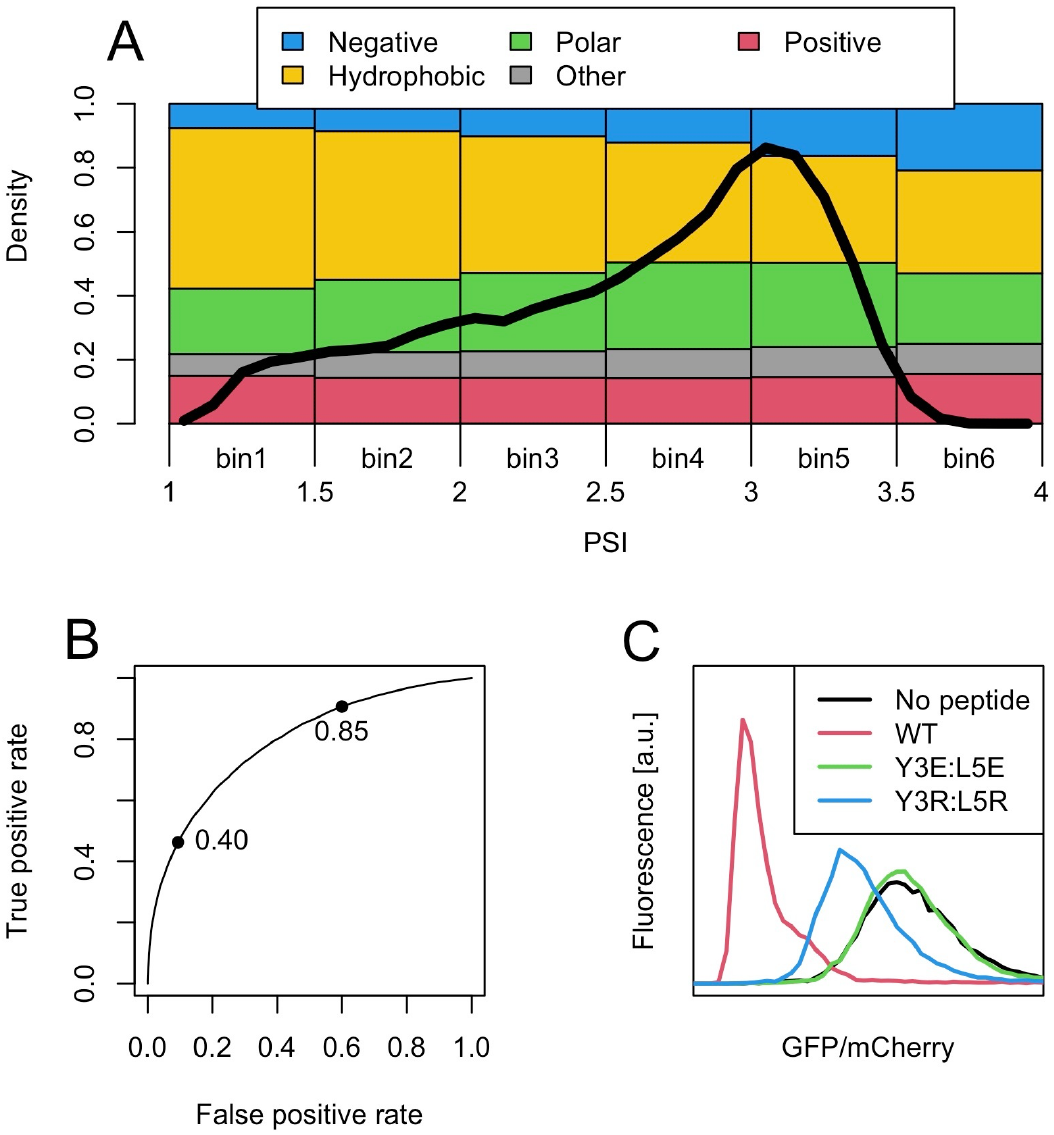
A) The amino acid compositions in six equally spaced PSI bins show a trend towards fewer hydrophobic (CAVMLIFYW, in yellow) and more negatively charged side chains (DE, in blue) in stable proteins (high PSI) compared to degrons (low PSI). Polar (NQST, in green), positively charged (HKR, in red) and other amino acids (GP, in grey) change less across the PSI distribution. The black line shows the distribution of PSI values for 18599 peptides (1175, 2406, 3355, 5599, 5877 and 187 peptides for bins 1 through 6 respectively). B) A logistic regression model that takes amino acid composition as input captures these trends well with an AUC of 0.79. The two points indicate that applying threshold values of 0.85 or 0.40 give a true-positive rate of 91% or a false positive rate of 9% respectively. C) A validation experiment where two degron potent amino acids (Tyr and Leu) in a degron tile (red) are substituted with Glu (green) or Arg (blue) to show the stabilizing effect of the changed composition.

We explore the predictive power of this observation using a logistic regression model that takes the amino acid composition as input (see Methods). This choice of model is motivated by a bimodal distribution of the PSI scores (Fig. 1A), which we simplify while building the prediction model by dividing peptides into those with degron activity (positive label) and those without (non-degrons or negative label). This model captures the trend well with an area under the curve (AUC) of 0.79 in a receiver operating characteristic (ROC) curve analysis. To further explore which part of the data is best described by this model, we trained three models that each excluded a slice of peptides with intermediate PSI. These showed an increasingly accurate fit to the data with an AUC up to 0.88 and without a decrease in the AUC when the models are evaluated on all data (Fig. S2). Thus, we conclude that peptides with intermediate PSI measurements do not contribute to the model accuracy and retrain a final model without these (Fig. 1B and Methods section). AUC values were retained in 10-fold cross validations. We term the final model QCDPred because the peptides were selected to represent the signals recognized by the PQC system ^18^. Python and R code to run QCDPred is available at https://github.com/KULL-Centre/papers/tree/main/2022/degron-predict-Johansson-et-al and a webserver is available at https://colab.research.google.com/github/KULL-Centre/papers/blob/main/2022/degron-predict-Johansson-et-al/QCDpred.ipynb.

While the AUC evaluations above suggest that much of the information in the experimental data is contained within the overall amino acid composition, we also explored whether a more complex model that use the di-peptide composition as input, could achieve a better fit. Each 17-residue peptide was thus represented by 16 dipeptides, with the idea that these would hold some context information for each amino acid. The accuracy of this model was, however, only slightly better with and AUC of 0.80 for the best model. Furthermore, the parameters of the dipeptide model (Fig. S3) were strongly correlated (Pearson correlation coefficient of 0.88) with the average of the two corresponding single amino acid parameters, suggesting that approximately the same information is contained in the simpler model (Fig. S4). The high accuracy of these amino acid composition models supports the observation that the ability of a peptide to act as a PQC degron is—to a large extent—determined by its amino acid composition and that hydrophobic peptides are strong degrons. The minor fraction of peptides that are not predicted accurately, as well as peptides of intermediate PSI, may indeed be recognised by more specific sequence or structural features; however, we do not attempt to separate such effects from noise in the measurements. We stress that we do not show that composition is the sole determinant of degradation, but rather that it has a substantial contribution and that we have not been able to find an additional signal from the data that we analysed.

The simple logistic regression model used in QCDPred has the advantage of being easily interpretable with the estimated parameters giving the degron contribution of the individual amino acid types—in the context of a 17 amino acid peptide (Fig. 2). The values in our model confirm our initial observation that the hydrophobic side chains in Met, Val, Leu, Ile and Cys, together with the aromatic side chains in Phe, Tyr and Trp, all promote degron activity. In contrast, the negatively charged side chains in Asp and Glu counteract degradation and have a strongly stabilising effect, whereas the positively charged side chains in Lys and—in particular Arg—have much smaller effects. Asn, Gln, Gly and Pro have intermediate stabilising effects. This interpretation and the generality of the observation is supported by results from related experiments in mammalian cells ^14^.

**Figure 2.**
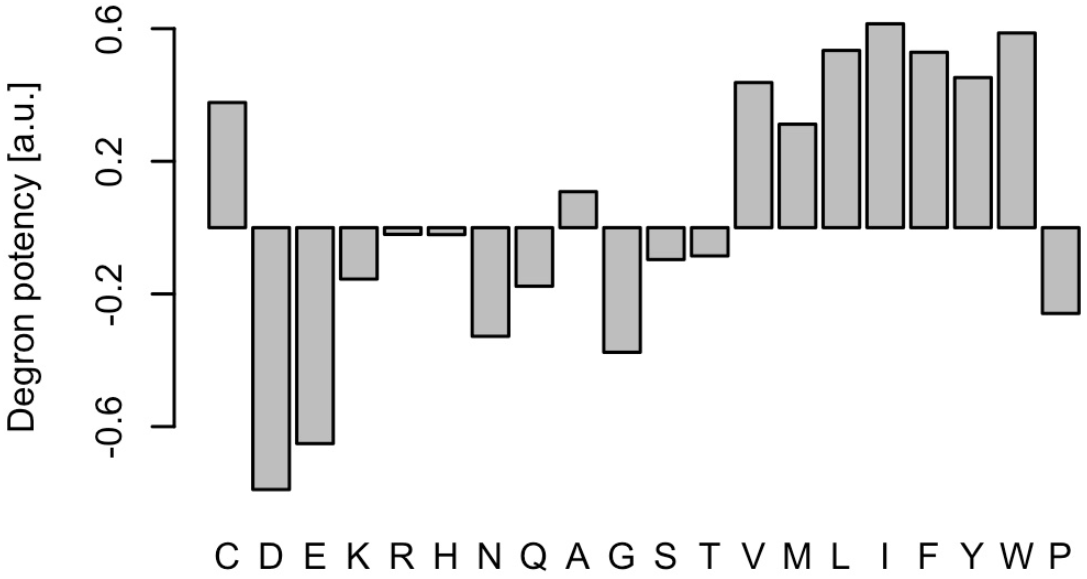
Estimated parameters for the logistic regression model may be interpreted as the degron propensity of individual amino acid types.

To validate the degron potency of individual amino acids, we selected a highly degron-active peptide (SLYLLSIWVKKFKWAGI) for additional analysis. In particular, we substituted two degradation promoting amino acids in this peptide, Tyr3 and Leu5, with either Arg or Glu. QCDPred predicts the wild-type peptide to be a strong degron and that the substitution of both Tyr and Leu with Glu to have a large effect on the degron capacity, and the substitution to Arg to be intermediate between wild-type and the Glu variant (supplementary Table S1). The degron effect of these peptides were evaluated using the same reporter system and flow cytometry readout as for the training data ^18^ (Fig. 1C), and the results show that the negative charges indeed have a stabilizing effect, and that the effect of the positively charged Arg lies between Glu and the wild-type Tyr and Leu.

Next, we examined whether the presence of degrons in proteins might be related to their cellular abundance. We consider 4533 proteins from the yeast proteome that exclude proteins containing any trans-membrane regions in this analysis because trans-membrane regions are enriched in hydrophobic amino acids that QCDPred would recognise, although these are unlikely to get exposed to the PQC system of the cytosolic environment ^19^. Indeed, as we show elsewhere aberrant exposure of trans-membrane regions to the PQC leads to degradation ^18^. The disordered regions, as assigned previously ^19^, of the remaining proteins are expected to be in contact with the PQC system whose effect is captured by QCDPred, and we thus hypothesized that the presence of degrons in these sequences would lead to decreased abundance. Indeed, we find that the average degron probability of these disordered residues correlates with protein abundance ^19, 20^ (Fig. 3a). While other mechanisms have also been suggested to lead to the observed relationship between amino acid composition and abundance ^19, 21^, our results suggest that QCDPred captures a significant degradation signal and thus a general mechanism of proteolytic stability in yeast cells. In line with this hypothesis that decreased abundance is due to more rapid degradation, we also find a correlation between QCDPred scores and protein half-life ^22^ (Fig. 3b). The signal is not very strong in the sense that a linear model of bin number only explains 10% and 15% of the total variance for abundance (bin>2) and half-life respectively, however, due to the high number of proteins included here, the results is highly significant, both with (nominal) p-values < 10^-15^ (based on an F-statistic but limited by a double precision floating point format). Comparisons across individual bins support this conclusion (supplementary Fig. S5).

**Figure 3.**
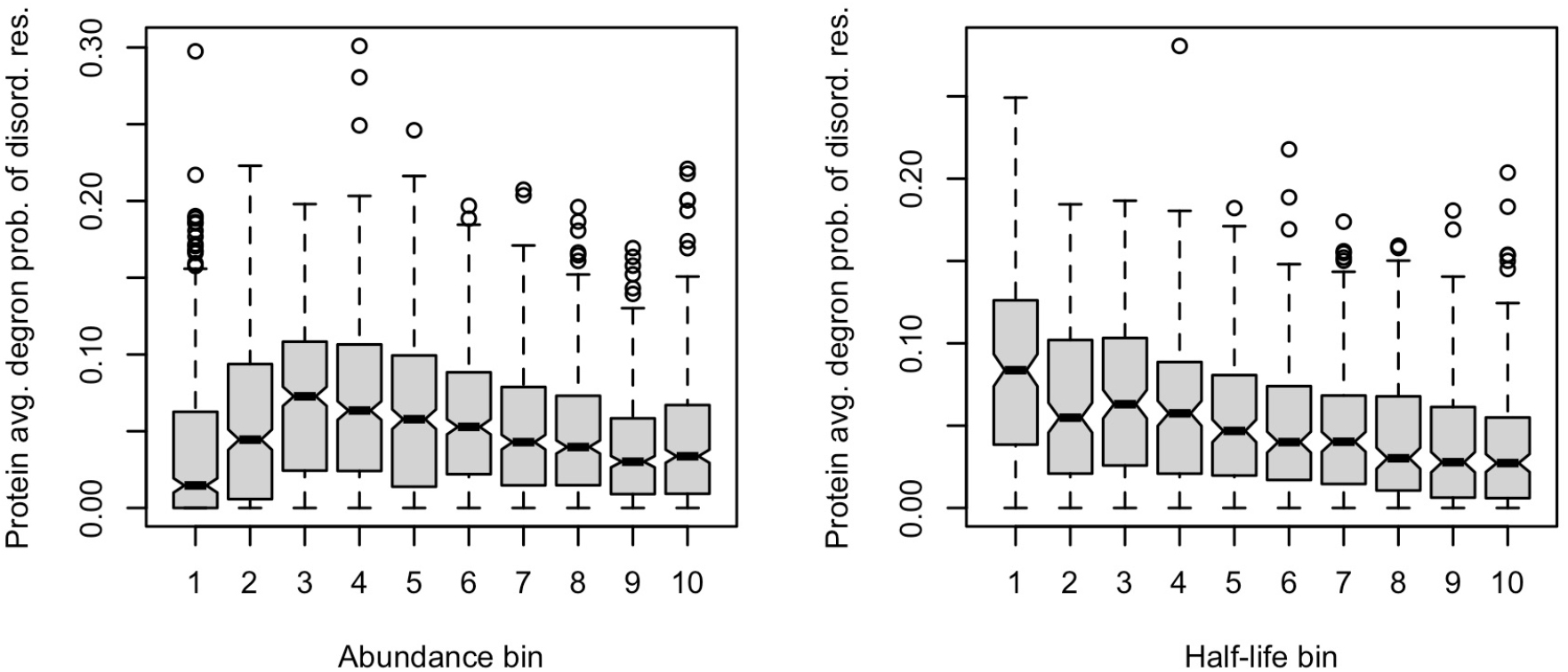
The average degron probability of disordered residues decreases for proteins of the yeast proteome that are more abundant and has longer half-life. Each bin contains approximately the same number of proteins; 445–462 proteins per abundance bin and 199– 201 proteins per half-life bin.

QCDPred has the advantage of being fast to evaluate, and we thus calculated the degron probability score for the 2.2 million residues in the 4533 yeast proteins in less than a minute (data available online at https://github.com/KULL-Centre/papers/tree/main/2022/degron-predict-Johansson-et-al). We confirm the QCDPred score distribution of our training data represents the 4533 proteins well (supplementary Fig. S6) and that both datasets have enrichment of degrons near the N-terminus compared to the C-terminus (supplementary Fig. S7). Similarly, ca. 24% of the peptides in our training data are predicted to be disordered (according to AlphaFold 2 pLDDT scores < 70)—in the context of the full-length proteins the sequences were obtained from—comparable to 31% for the AlphaFold 2 models of 6040 yeast proteins ^23, 24^.

We note that the average degron probabilities of disordered regions are in general low, which is expected since strong degradation signals in these regions should cause rapid degradation (unless, for example, that the sequences are buried in complexes inside the cell). We also note that in our analysis of protein abundance the two lowest-abundance bins are exceptions to the overall correlation which might be explained by the low concentrations of these proteins. The protein concentration corresponds to approximately 5-30 nM which is lower than many typical binding affinities ^19^, thus perhaps bypassing the degradation mechanism.

## Conclusions

Here we present the—to our knowledge—first prediction model for PQC degrons. This work complements previous work on mapping regulatory degrons, which are typically more specific. Our results confirm previous findings that hydrophobic sequences are recognized and targeted for degradation ^14, 25, 26^. We also find clear differences between how different polar and charged residues contribute to degrons. Most notably, we find that the presence of the negatively charged side chains in Asp and Glu counteract degradation whereas the positively charged side chains in Lys and Arg have a more neutral effect. These results complement our recent observation that Hsp70 binding motifs, which are often hydrophobic and positively charged, can act as degrons ^17, 27^. The effects of hydrophobic, positively and negatively charged residues are also validated by experiments that show that replacing hydrophobic residues with Glu in a degron can prevent degradation, whereas replacing the same amino acids with Arg has a substantially smaller effect^18^. We stress that while QCDPred is only based on sequence composition this does not mean that the actual amino acid sequence is not important for PQC degrons. Instead, our results imply that composition plays a substantial role, and we suggest that larger and different libraries will help determine the role of the actual sequence. Indeed, scrambling the sequence of a degron peptide can affect its strength ^18^.

In the future, it will be interesting to explore the effects of degron properties further in the context of cellular functions and evolution. Here we note for example that transactivation domains of transcription factors are enriched in hydrophobic and negatively charged residues that are known to be key for function ^28, 29^. We speculate that the presence and necessity of the negatively charged residues in these may not only be due to direct effects on function, but also to counteract the higher degradation propensity afforded by the presence of for example many aromatic residues, thus suggesting that the molecular evolution of unstructured regions of transcription factors have been shaped by the PQC system. In addition, it has been found that negatively charged proteins only rarely engage with molecular chaperones ^30–32^. More generally, we show a clear relationship between the presence of degrons in disordered regions and the abundance and degradation rates of yeast proteins. In this context we note that human disease mutations in disordered regions are substantially enriched in Glu-to-Lys substitutions, as well as Arg-to-Trp and Arg-to-Cys substitutions ^33, 34^, that we suggest could lead to more rapid degradation of these proteins. Finally, we note that the sequence properties that we have mapped also resemble those of transmembrane regions, and indeed we elsewhere show that when transmembrane regions are mislocalized to the cytosol they are also targeted for degradation ^18^.

In many cases, the effects discussed above can be traced back to differences in hydrophobic content of the amino acid sequences. For example, the overall amino acid compositions of disordered and folded proteins are different ^35^, and the depletion of hydrophobic amino acids in disordered proteins underlies their inability to fold into specific three-dimensional structures. Thus, by recognizing hydrophobic sequences, the PQC system can distinguish misfolded globular or transmembrane proteins from for example disordered regions or those on surfaces of folded proteins, and in this way display functional specificity without substantial sequence specificity.

More than half of disease-causing missense variants are thought to lead to protein degradation ^36^, but the molecular mechanisms of how these variants are recognized by the cell is still incompletely understood. Our simple, but accurate prediction algorithm to identify degrons from sequence may help understand the complex relationship between sequence variation and abundance. We and others have previously shown that many disease-causing missense variants are thermodynamically destabilized ^27, 37–43^. The results presented here provide a key missing part of the puzzle, namely a quantitative model for which exposed sequences are PQC targets, and we hope our work may help develop quantitative models that combine protein stability and recognition via the PQC.

While our work is a substantial step forward, there are some limitations. First, in the library we have screened we only vary 17-residue long peptides, and so our model only predicts degrons that fit within this length. More importantly, an unfolded or misfolded protein may have multiple degrons and we have previously shown that an increased number of exposed degrons lead to progressively lower abundance ^17^. Thus, future work should quantify the relationship between the length and number of degrons and protein abundance. Our model also does not provide additional insight into the presence of potentially specific sequence motifs. We believe this is mostly due to two, related effects. First, the PQC system needs to be able to target a wide range of proteins, and thus the PQC may have evolved to recognize generic properties of misfolded proteins such as exposed hydrophobic cores. Indeed, the yeast PQC E3 San1 appears to broadly recognize exposed hydrophobicity rather than specific sequence motifs ^25, 44^. Other PQC E3s such as Ubr1 ^45^ and STUB1/CHIP ^46^ have relegated substrate recognition to molecular chaperones, who also display promiscuity in their substrate selection ^47, 48^. Second, there is likely extensive redundancy in the components of the PQC system including E3 enzymes, chaperones and adaptor proteins ^16, 26, 49–51^. Thus, a more extensive set of experiments using different genetic perturbations are likely needed to disentangle the sequence preferences of different components. Finally, we note that our experiments and analyses are focused on detecting linear degrons. While we expect that these are important in the general PQC system, future experiments and computational analyses are needed to discover potential PQC degrons that are perhaps localized in a three-dimensional structure but not in the linear polypeptide chain.

In summary, we conclude that the approach we have developed provides a first, key step in understanding and predicting the sequence properties of PQC degrons. We hope that the computational model, QCDPred, the experimental data ^18^ and the application to the yeast proteome will be useful to understand better the determinants of protein abundance and quality control.

## Methods

### Logistic regression model

The data used for training the model is described in ^18^. Briefly, the library consists of 326 proteins from 24 yeast complexes, all tiled into peptides of 17 amino acids with 12 amino acids overlap. For each peptide, a protein stability index (PSI) is calculated as ∑_*g*∈{1.4}_ *g* × *p_i,g_* where *p_i,g_* is the population of peptide *i* in each of the four FACS gates, *g*. The PSI values thus range from one (most degron prone) to four (most stable) and reflects the average position among the four quartiles of the FACS distribution. In this work, we consider the 18,599 most confident peptide measurements with more than 50 sequencing reads across all gates, representing 323 proteins with a coverage of 93% of residues. Of these peptides, 4790 with PSI < 2.2 were labelled as degron-active (positive) and 8769 peptides with PSI > 2.8 as degron inactive or stable (negative). The degrons are diverse and represents 296 of the 323 assayed proteins. Based on a 10-fold cross validation, the strength of the ridge regularization penalty in the logistic model was set to 0.001 and the model was retrained on all the labelled data resulting in the same AUC of 0.85 as during cross validation. To evaluate the model on data that includes unlabelled peptides with intermediate PSI, we labelled these as degrons if PSI < 2.5 and non-degrons otherwise. The same procedure was followed for the dipeptide model using a regularization strength of 1.0. Models were made in *The R project for statistical computing* [R] ^52^ using the *glmnet* package ^53^. Code and data are available at https://github.com/KULL-Centre/papers/tree/main/2022/degron-predict-Johansson-et-al.

### Validation experiment

The DNA sequences encoding the validation peptide R5 and its mutants (see supplementary Table S1 for details) were re-cloned into the parental vector and were tested and analyzed as described ^18^. Briefly, cells were grown on selective media to mid-log-phase and subjected to a CellStream analyzer instrument (Merck) equipped with 488 nm and 561 nm lasers for detecting GFP and mCherry, respectively. 10,000 events of each peptide were plotted against the ratio of GFP to mCherry.

### Abundance and half-life analysis

For the half-life and abundance analyses, we used a collection of 4533 proteins from Dubreuil *et al.* ^19^ annotated with cellular abundance levels from the Pax database ^20^ and residues predicted to be disordered by IUPRED ^54^. The 30% of residues with highest IUPRED score were assumed to be disordered. This set is made by excluding proteins with any trans-membrane regions from the yeast proteome and is particularly relevant here because trans-membrane regions are expected to be highly hydrophobic. For 1993 of the 4533 proteins in this set we were able to assign a half-life ^22^. For each protein we calculated the QCDPred degron probability of each consecutive 17-mer and assigned the value to the central residue position. Finally, for each protein we averaged the degron probability of the positions that are predicted to be disordered. These predictions are available at https://github.com/KULL-Centre/papers/tree/main/2022/degron-predict-Johansson-et-al and predictions for the full yeast proteome are available at https://erda.ku.dk/archives/3f5bf0ada384be330541a3c1b04168d9/uniprot_2022_01_proteome_UP000002311_qcdpred.txt.gz.

## Acknowledgments

This work is a contribution from the Novo Nordisk Foundation centre PRISM (NNF18OC0033950; to R.H.-P. and K.L.-L.), and also acknowledges support from a NSF-BSF grant (2016722 ; to T.R). B.M. acknowledges the support of the Neubauer doctoral fellowship fund.

## Abbreviations

PQC: Protein quality control
PSI: Protein stability index
ROC: Receiver operating characteristic
AUC: Area under the curve

