## Supporting Figures and Table for "Prediction of quality-control degradation signals in yeast proteins"

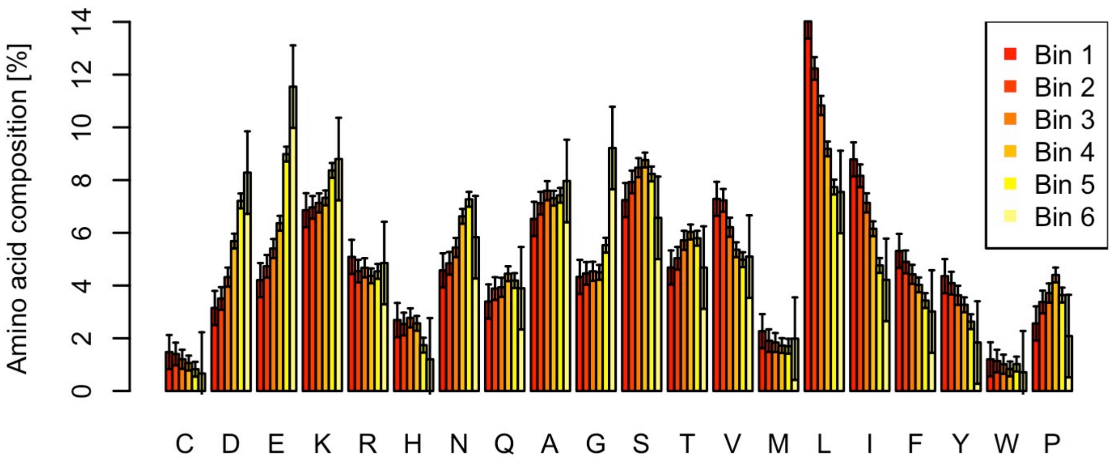


**Figure S1.** Composition of each amino acid in the 6 bins across the PSI distribution (Fig. 1).

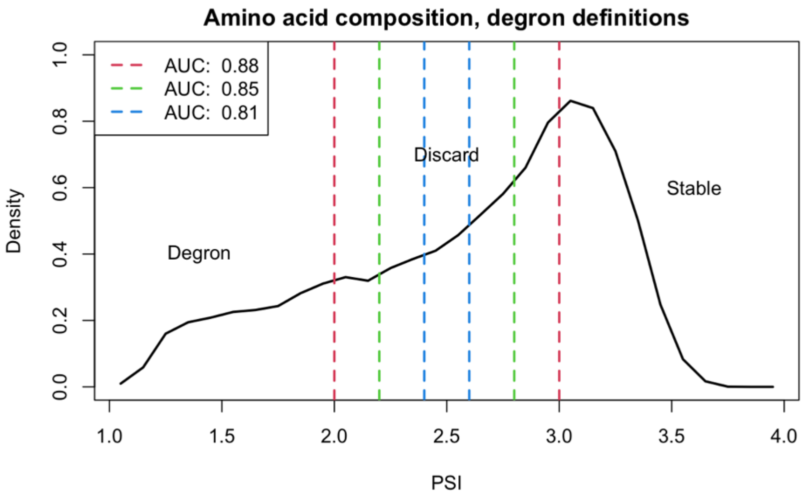


***Figure S2.*** *Peptides with intermediate PSI values do not contribute to model accuracy when included in training. We trained three models in which we excluded different sets of peptides with intermediate PSI values. The AUCs of the resulting models increase as more peptides of intermediate PSI are excluded. When we evaluate these three models on the full dataset (that is when also including the intermediate PSI values) we obtain an AUC of 0.79 for all three models.*

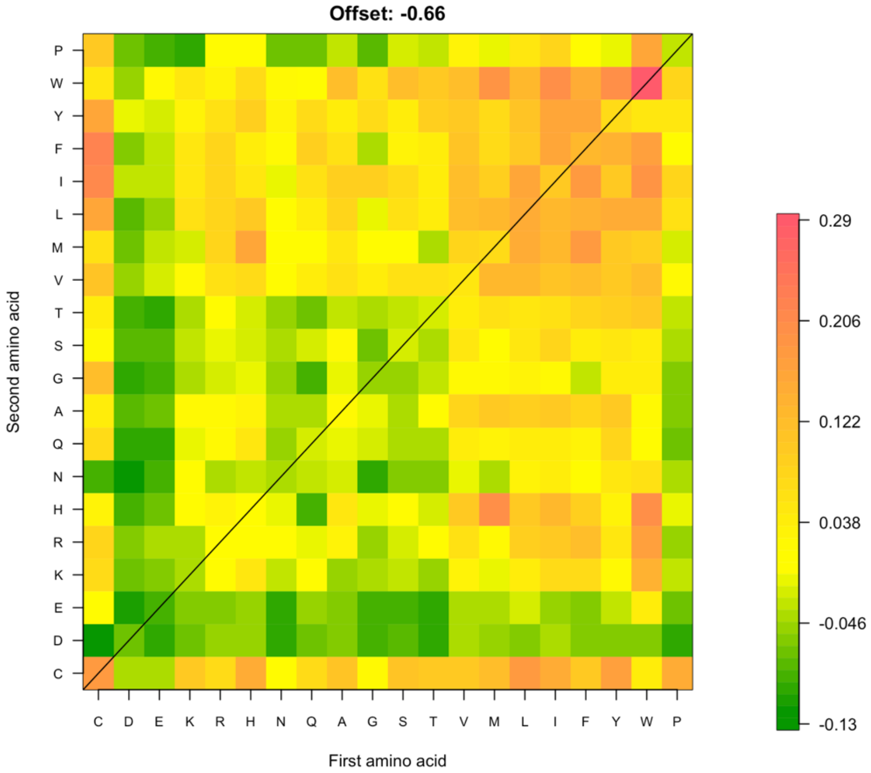


***Figure S3.*** *Estimated parameters for a logistic regression model that uses di-peptide composition as input.*

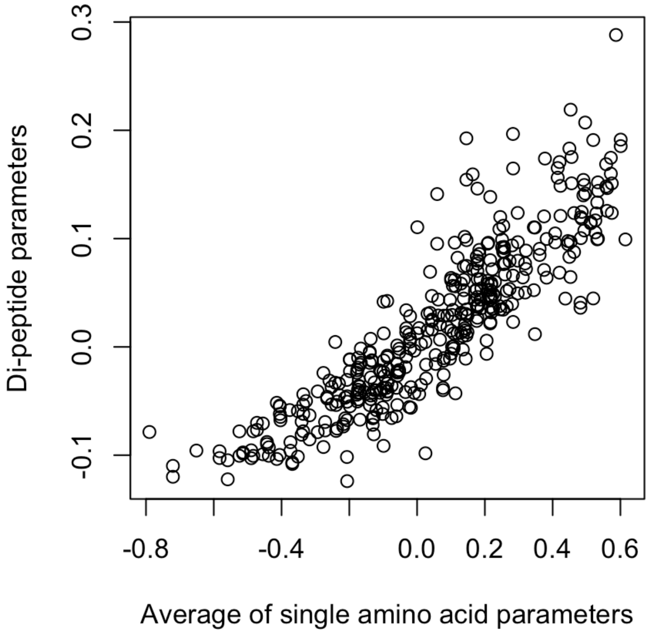


***Figure S4***. *The parameters estimated parameters in the dipeptide model correlate strongly (Pearson correlation coefficient 0.88) with the average of the two corresponding single amino acid parameters of the single amino acid composition model.*

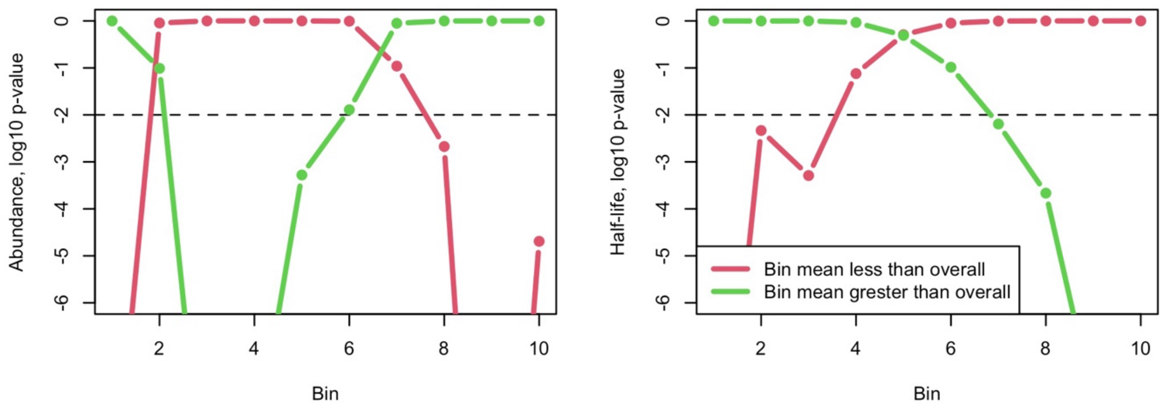


***Figure S5.*** *Welch two-sample t-test for average degron probability of individual bins in Fig. 3 being less than (red) or greater than (green) the overall average degron probability. Bins 3,4,5,8,9 and 10 for abundance (left) and bins 1,2,3,7,8,9 and 10 for protein half-life (right) may be considered significantly different from the overall average ($p < 0.01$).*

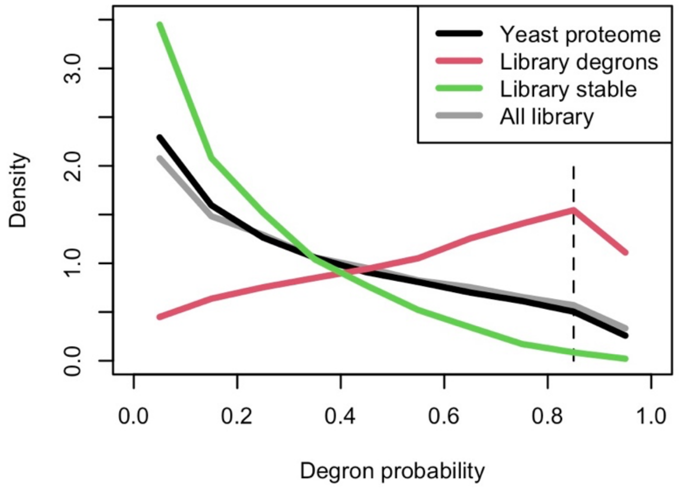


***Figure S6.*** *Distribution of QCDpred scores for the tiled trans-membrane free yeast proteome (black line; 433,738 tiles) compared to all library tiles (grey line; 18,599), training data labelled as degrons (red line; 4,790 tiles) or stable (green line; 8,769 tiles). The proteome behaves approximately like our library. The slight enrichment of stable tiles in the proteome may reflect a higher fraction of disordered protein. The proteome proteins are tiled in the same way as the library for this analysis.*

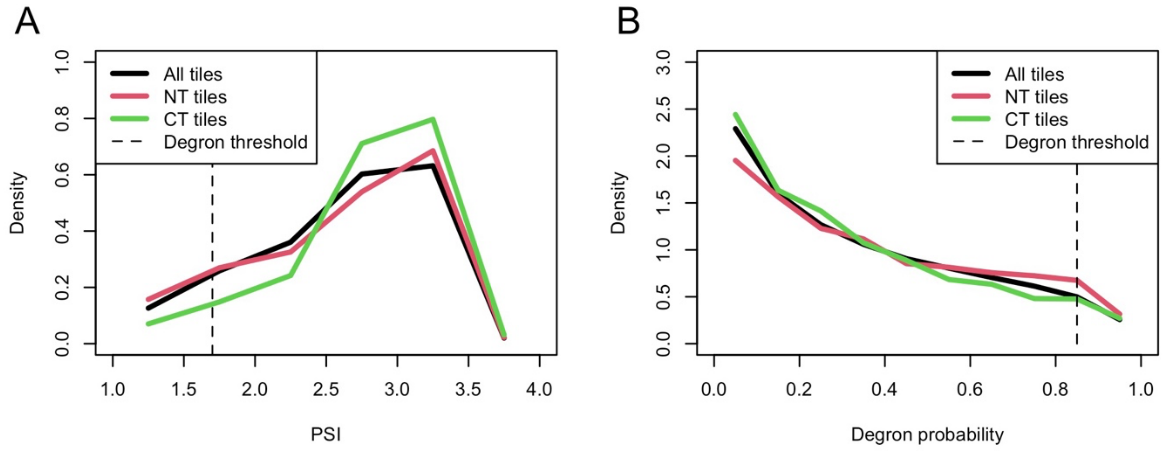


***Figure S7.*** *Asymmetry of degrons near the N- and C-termini. A) PSI distribution of the training data compared to the N-terminal (red) and C-terminal tiles (green). The fraction of degron active tiles (PSI < 1.7; dashed line) is 10.9%, 12.4% and 5.5% for 18,616 total, 178 N-terminal and 256 C-terminal tiles respectively. B) QCDpred score distribution of the trans-membrane free yeast proteome (black line) compared to N-terminal tiles (red line) or C-terminal tiles (green line). The fraction of degron-prone positions (tile score >= 0.85) is 4.9% for all tiles, 6.5±0.3% for N-terminal tiles, and 4.3±0.3% for C-terminal tiles. Bootstrap uncertainties are calculated from resampling the 4533 terminal tiles with replacement 1000 times.*

| Peptide | Sequence | Gene | Start | End | QCDPred score |
|---|---|---|---|---|---|
| Y3E:L5E | CCGGGTCTTTGgagTTGgagTCTATCTGGG TGAAGAAGTTCAAATGGGCCGGTgagTGAG | CYT1 | 279 | 295 | 0.51 |
| WT | CCGGGTCTTTGTATTTGCTATCTATCTGGG TGAAGAAGTTCAAATGGGCCGGTATCTGAG | CYT1 | 279 | 295 | 0.97 |
| Y3R:L5R | CCGGGTCTTTGcgtTTGcgtTCTATCTGGG TGAAGAAGTTCAAATGGGCCGGTcgtTGAG | CYT1 | 279 | 295 | 0.87 |

***Table S1.*** *Peptides used in the validation experiments.*
